# DPPA2 and DPPA4 are dispensable for mouse zygotic genome activation and preimplantation development

**DOI:** 10.1101/2021.08.19.457017

**Authors:** Zhiyuan Chen, Zhenfei Xie, Yi Zhang

## Abstract

How maternal factors in oocytes initiate zygotic genome activation (ZGA) remains elusive. Recent studies indicate that DPPA2 and DPPA4 are required for establishing a 2-cell embryo-like (2C-like) state in mouse embryonic stem cells (ESCs) in a DUX-dependent manner. These results suggest that DPPA2 and DPPA4 are essential maternal factors that regulate *Dux* and ZGA in embryos. By analyzing maternal knockout and maternal-zygotic knockout embryos, we unexpectedly found that *Dux* activation, ZGA, and preimplantation development are normal in embryos without DPPA2 or DPPA4. Thus, unlike in ESCs/2C-like cells, DPPA2 and DPPA4 are dispensable for ZGA and preimplantation development.

## Introduction

Following fertilization, embryonic development relies on maternal factors deposited during oogenesis initially and then the newly generated embryonic product after its genome is activated. The awakening of embryonic genome is known as zygotic or embryonic genome activation (ZGA/EGA), which is fundamental for an embryo to acquire totipotency and to undergo normal development. In mice, ZGA is consisted of two successive waves of transcription with a minor and a major wave occurring at late 1-cell and late 2-cell stage, respectively (Schultz et al. 2018). Transcription at late 1-cell stage is largely promiscuous and the transcripts are typically inefficient of splicing and 3’processing (Abe et al. 2015), whereas the major ZGA at late 2-cell stage are coupled with the expression of thousands of translatable mRNAs (Hamatani et al. 2004; Wang et al. 2004; Zeng et al. 2004). Notably, both minor and major ZGA are essential for mouse preimplantation development (Abe et al. 2018; Schultz et al. 2018; Liu et al. 2020).

The 2C-like cells are a rare cell subpopulation in ESCs that is characterized by the expression of many 2C-specific transcripts such as *Zscan4* and *Zfp352* and has the expanded potential to contribute to both embryonic and extraembryonic lineages (Macfarlan et al. 2012). The 2C-like cells have been widely used as an *in vitro* approximate for understanding totipotency and ZGA (Fu et al. 2020; Genet and Torres-Padilla 2020). Studies in the past several years have revealed DUX (human homologue as DUX4), a double homeodomain protein, as a master regulator of 2C-like state in ESCs (De Iaco et al. 2017; Hendrickson et al. 2017; Whiddon et al. 2017). Most of the mechanisms that promote 2C-like state in ESCs identified so far directly or indirectly regulate *Dux* activation. Intriguingly, loss of *Dux* in embryos only mildly affects ZGA and the *Dux* null embryos are viable with reduced litter sizes (Chen and Zhang 2019; Guo et al. 2019; De Iaco et al. 2020; Bosnakovski et al. 2021). Thus, although DUX is important for synchronizing and enhancing the expression of some 2C embryo-specific genes, it is not essential for ZGA and embryogenesis.

Since *Dux* is not expressed in oocytes and it gets activated only at late 1-cell stage, upstream factors should have already existed in oocytes to trigger *Dux* expression during minor ZGA. Recent studies have identified the developmental pluripotency associated 2 and 4 (*Dppa2* and *4*) as essential factors for establishing the 2C-like state in ESCs by activating *Dux* (De Iaco et al. 2019; Eckersley-Maslin et al. 2019; Yan et al. 2019). In addition, DPPA2 and DPPA4 directly regulate young LINE-1 elements in ESCs in a DUX-independent manner (De Iaco et al. 2019). The young LINE-1 elements levels also increase during the major ZGA. Taken together, these findings suggest that *Dppa2 and Dppa4*, which are expressed in oocytes, may regulate ZGA as maternal factors through both DUX-dependent and -independent pathways. In support of this, overexpression of the dominant negative forms of *Dppa2* has been shown to impair mouse preimplantation development (Hu et al. 2010; Yan et al. 2019).

Despite the above evidence, whether oocyte derived DPPA2 and DPPA4 activate *Dux* expression and regulate ZGA in mouse embryos has not been rigorously examined. The previous *Dppa2* and *Dppa4* zygotic knockout (KO) studies (Madan et al. 2009; Nakamura et al. 2011) did not address the maternal contributions of these two proteins in ZGA and preimplantation development. In this study, we generated maternal KO and maternal-zygotic KO mouse embryos for both *Dppa2* and *Dppa4*, and determined their functions in *Dux* activation, ZGA, and preimplantation development. Our results demonstrate that both Dppa2 and Dppa4 are dispensable for ZGA and preimplantation development.

## Results & Discussion

### Expression dynamics of *Dppa2 and Dppa4* in mouse early development

To test the possibility that DPPA2 and DPPA4 are involved in activating *Dux* expression and ZGA, we first determined the expression dynamics and cellular localization of these proteins in early embryos by RNA sequencing (RNA-seq) and immunostaining analyses. We found that low levels of *Dppa2* and *Dppa4* RNAs were detectable in both oocytes and zygotes, and their expression levels were dramatically increased during major ZGA and reached at peak at 8-cell stage **(Fig. 1A).** However, both *Dppa2* and *Dppa4* RNA became undetectable soon after embryo implantation **(Fig. 1A)** and their transcriptional silencing is presumably achieved by gain of DNA methylation at the promoters (Eckersley-Maslin et al. 2019).

**Figure 1.**
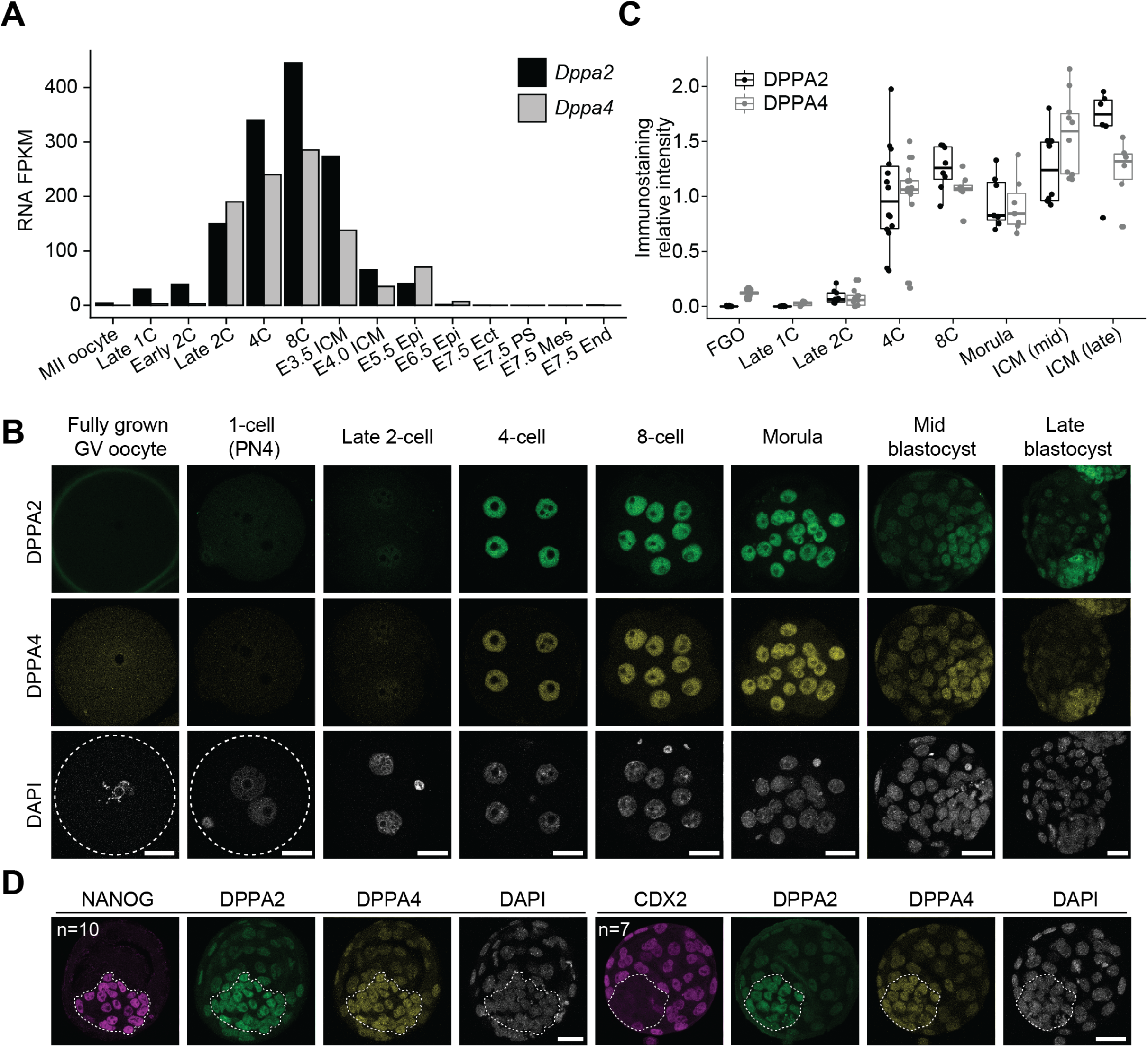
Expression and cellular localization of DPPA2 and DPPA4 in mouse oocytes and early embryos. **A)** RNA levels of *Dppa2* and *Dppa4* in oocyte and early embryos. The RNA-seq data were from (Wu et al. 2016; Zhang et al. 2018). **FPKM**: fragments per kilobase of transcript per million mapped reads; **E**: embryonic day; **ICM**: inner cell mass; **Epi**: epiblast; **Ect**: ectoderm; **End**: endoderm; **Mes**: mesoderm; **PS**: primitive streak. **B)** Images of oocytes and preimplantation embryos immunostained with antibodies against DPPA2 and DPPA4. **Scale bar**: 20 μm; **GV:** germinal vesicle; **PN**: pronucleus. **C)** Quantification of the signal intensities of DPPA2 and DPPA4. The average signal intensities of 4-cell embryos were set as 1.0. The total number of oocytes/embryos analyzed were 13 for oocytes, eight for 1-cell embryos, nine for 2-cell embryos, 14 for 4-cell embryos, eight for 8-cell embryos, seven for morulae, 10 for mid blastocysts, and six for late blastocysts, respectively. The middle lines represent medians. The box hinges indicate the 25^th^ and 75^th^ percentiles, and the whiskers indicate the hinger ± 1.5×interequatile range. **FGO:** fully grown GV oocytes. **D)** Images of blastocysts immunostained with antibodies against DPPA2, DPPA4, NANOG, and/or CDX2. Number of embryos analyzed are as labeled. **Scale bar**: 20 μm.

In contrast to the RNA levels, DPPA2 and DPPA4 immunostaining signals were not detectable in both oocytes and zygotes **(Fig. 1B and C)**. The signals were first detectable at late 2-cell and became stronger at the subsequent stages **(Fig. 1B and C)**. At blastocyst stage, DPPA2 and DPPA4 were mostly located in inner cell mass (*i.e.*, NANOG-positive) rather than trophectoderm (*i.e.*, CDX2-positive cells) (**Fig. 1D**). This observation is consistent with the previous RNA in situ hybridization experiments showing that *Dppa2* and *Dppa4* are restricted to inner cell mass at this stage (Maldonado-Saldivia et al. 2007).

### Generation of *Dppa2* and *Dppa4* maternal and maternal-zygotic KO embryos

The immunostaining results are incompatible with a potential role of DPPA2 and DPPA4 in *Dux* activation and ZGA in embryos. However, it is also possible that the very low levels of maternal DPPA2 and DPPA4 proteins in oocytes (barely detected by immunostaining) may still play a role in activating *Dux* and ZGA. To test this possibility, we generated *Gdf9-Cre-*mediated oocyte-specific conditional KO (CKO) models for *Dppa2* (exon 3-4 floxed) (**Fig. 2A**) and *Dppa4* (exon 2-7 floxed) (**Fig. 2B**). The *Gdf9-Cre* is expressed in early growing oocytes around postnatal day 3 (Lan et al. 2004). Since the *Dppa4* flox *(fl)* allele has been previously established and described (Nakamura et al. 2011), we only characterized in detail of the *Dppa2 fl* allele that was generated in this study using the 2-cell homologous recombination (2C-HR)-CRISPR method (Gu et al. 2018)(**Fig. S1A-B**). Sanger sequencing analyses confirmed that exon 3-4 of *Dppa2* were successfully depleted in the CKO oocytes (**Fig. S1C**), resulting in a frameshift with the disruption of both the SAP and C-terminal domains. Since *Dppa2* and *Dppa4* are closely linked on the same chromosome, it is not feasible to generate *Dppa2* and *Dppa4* double CKO mice by natural mating.

**Figure 2.**
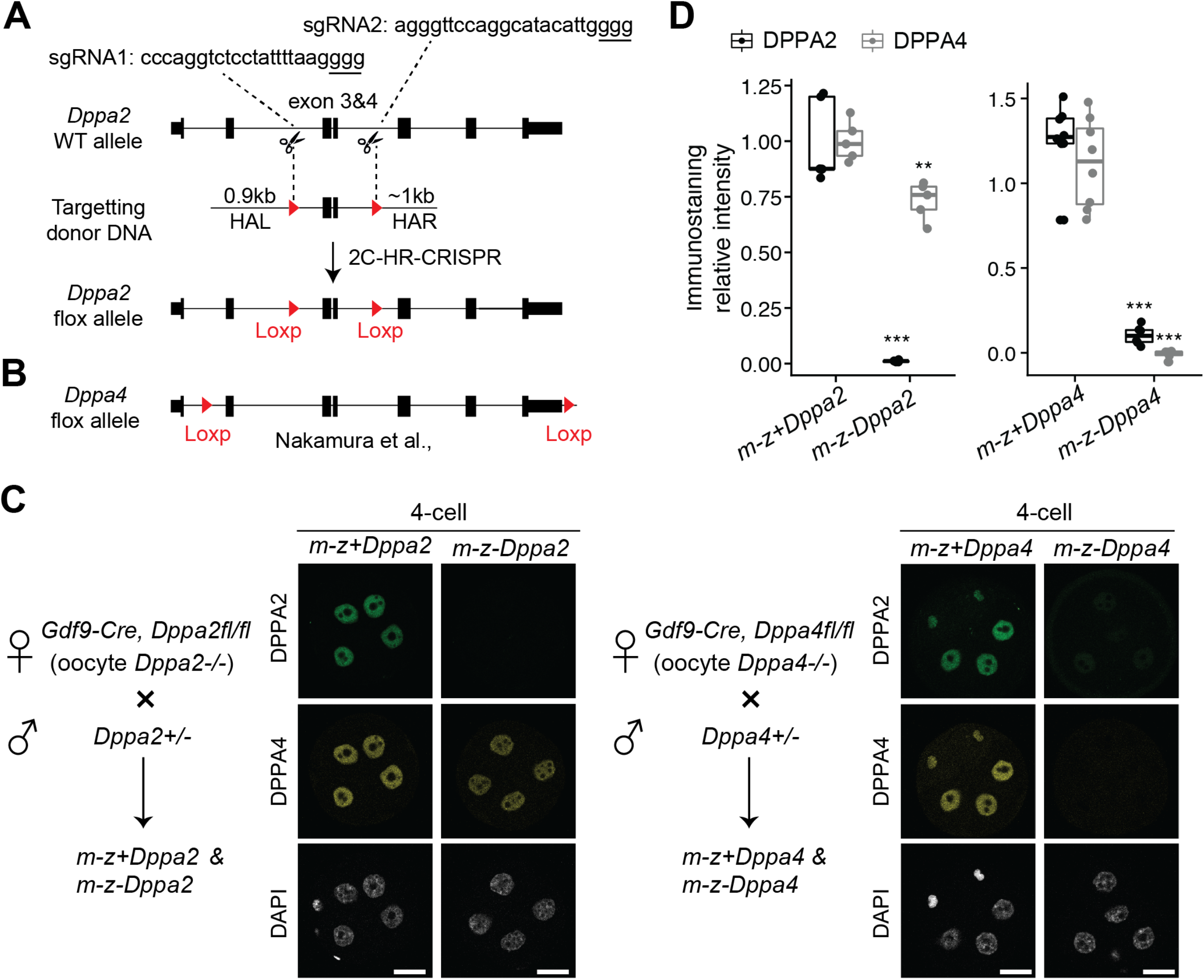
Generation of maternal KO (*m-z+*) and maternal-zygotic KO (*m-z−*) embryos for *Dppa2* and *Dppa4*. **A)** Gene targeting strategy for *Dppa2* flox allele. **HAL**: left homologous arm; **HAR**: right homologous arm; **HR**: homologous recombination. **B)** Schematics for *Dppa4* flox allele (Nakamura et al. 2011). **C)** Schematics for generating *m-z+* and *m-z−* embryos and images of 4-cell embryos immunostained with antibodies against DPPA2 and DPPA4. **D)** Quantifications of signal intensities of DPPA2 and DPPA4. The average signal intensities of *m-z+* embryos were set as 1.0. The number of embryos analyzed were five for *m-z+Dppa2*, five for *m-z−Dppa2*, eight for *m-z+Dppa4*, and six for *m-z−Dppa4*, respectively. The middle lines represent medians. The box hinges indicate the 25^th^ and 75^th^ percentiles, and the whiskers indicate the hinger ± 1.5×interequatile range. (***) *P* < 0.001, (**) *P* < 0.01, two-tailed Student *t-*test.

We next sought to confirm successful KO of DPPA2 and DPPA4 at the protein level. Because of the weak immunostaining signals of DPPA2 and DPPA4 before 4-cell stage (**Fig. 1B**), it was challenging to determine their signal loss in CKO oocytes and maternal KO 1-cell/2-cell embryos. To circumvent this issue, immunostaining analyses were performed at 4-cell stage for maternal KO (*m-z+*) and maternal-zygotic KO (*m-z−*) embryos generated by crossing CKO female mice with heterozygous male mice (**Fig. 2C**). The signal intensities of DPPA2 and DPPA4 in maternal KO 4-cells were largely comparable to the wild-type (WT) embryos (**Fig. 2C and Fig. 1B**). This suggests that *Dppa2* and *Dppa4* were zygotically expressed from the WT paternal alleles, which compensated for the maternal loss. In contrast, DPPA2 and DPPA4 signals were not detectable in *m-z−Dppa2* and *m-z−Dppa4* 4-cell embryos, respectively (**Fig. 2C-D**), confirming the successful KO of these proteins. It is worth noting that loss of one protein caused reduced signal of the other in maternal-zygotic KO 4-cell embryos, albeit to different extents (**Fig. 2C-D**). This is perhaps due to the fact that DPPA2 and DPPA4 function as a heterodimer (Nakamura et al. 2011; Hernandez et al. 2018) and loss of one affected the stability of the other as has been observed in ESCs (Gretarsson and Hackett 2020).

### DPPA2 and DPPA4 are not required for preimplantation development

Having confirmed the successful KO of DPPA2 and DPPA4 in embryos, we next examined the KO impact on preimplantation development. To this end, spermatozoa from WT male mice were used to fertilize oocytes from control and CKO female mice *in vitro*, generating WT (*m+z+*) and maternal KO (*m-z+*) embryos. To exclude the possibility that WT paternal copy may compensate for the maternal losses, CKO oocytes were also fertilized with heterozygous spermatozoa, which should generate maternal-zygotic KO (*m-z−*) in half of the embryos. Unexpectedly, none of the embryo groups showed apparent preimplantation defects (**Fig. 3A-B**). Consistently, the maternal-zygotic KO embryos for DPPA2 and DPPA4 can develop to blastocyst with the formation of pluripotent NANOG-positive inner cell mass (**Fig. 3C**). Notably, loss of both DPPA2 and DPPA4 in *m-z−Dppa4* blastocysts making these embryos equivalent to *Dppa2/4* double KO (**Fig. 3C**). These results indicate that DPPA2 and DPPA4 are not required for mouse preimplantation development.

**Figure3.**
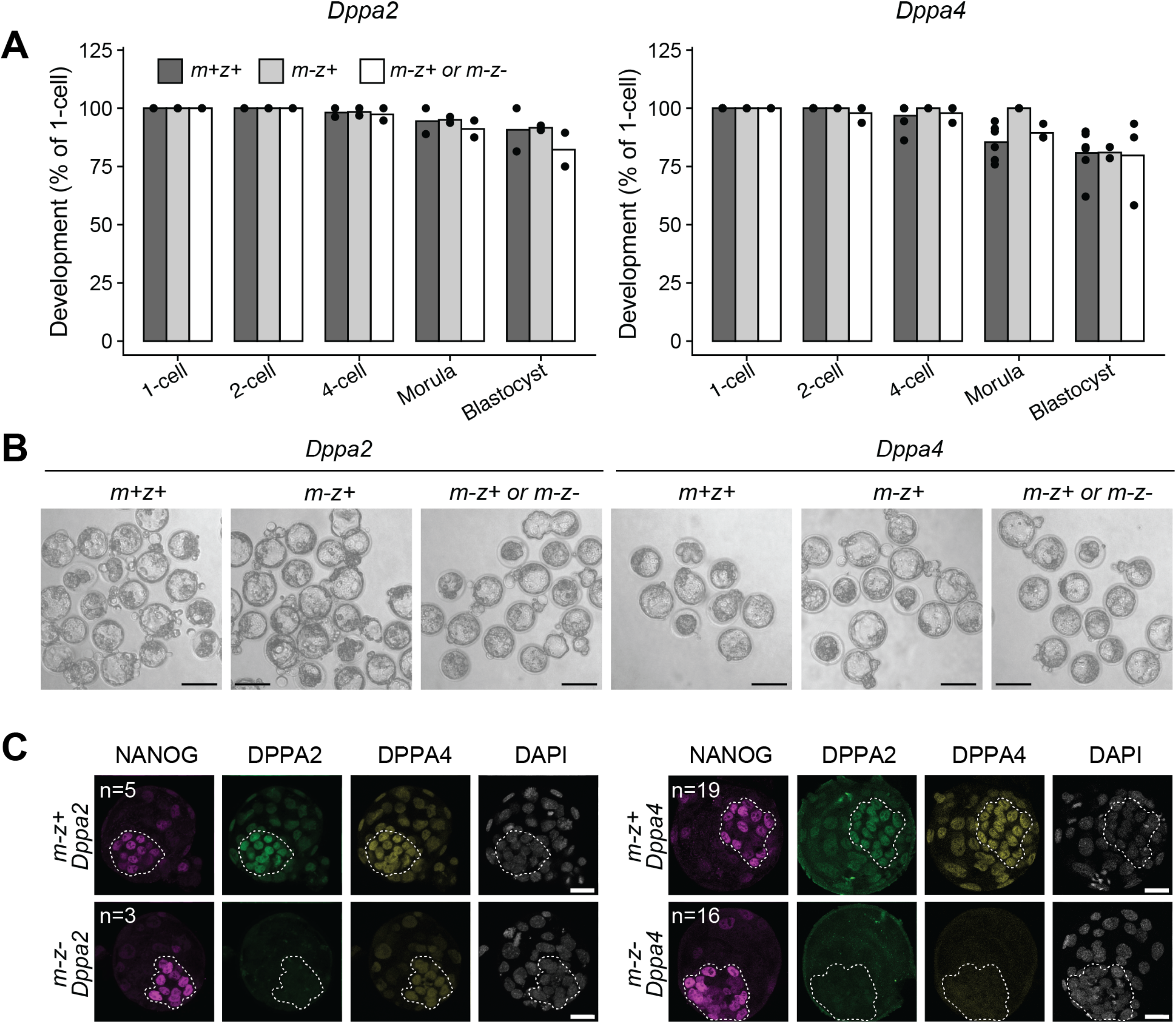
Embryos without DPPA2 and DPPA4 undergo normal preimplantation development. **A)** Bar graphs showing the percentage of embryos reaching the indicated developmental stages. The number of experiments performed are denoted by dots. Total number of embryos analyzed were 44 for *m+z+Dppa2*, 59 for *m-z+Dppa2*, 27 for *m-z+/m-z−Dppa2*, 95 for *m+z+Dppa4*, 20 for *m-z+Dppa4*, and 50 for *m-z+/m-z−Dppa4*, respectively. **B)** Images of embryos after 96 hours of culturing *in vitro*. **Scale bar**: 80 μm. **C)** Images of blastocysts immunostained with antibodies against NANOG, DPPA2, and DPPA4. The embryos were generated by fertilizing CKO oocytes with heterozygous spermatozoa. Number of embryos analyzed are as labeled. **Scale bar**: 20 μm.

### DPPA2 and DPPA4 are dispensable for *Dux* expression and ZGA

Given that mouse preimplantation development is largely normal without DPPA2 or DPPA4, minimal ZGA defects are expected in these mutants. To confirm this prediction, we performed RNA-seq experiments. To determine whether DPPA2 and DPPA4 initiate *Dux* transcription during minor ZGA and subsequently affect major ZGA, late 1-cell and late 2-cell embryos of control and maternal KO were collected for RNA-seq analyses (**Fig. S2A**). Maternal-zygotic KO single 2-cell embryos were also analyzed to exclude the possibility that WT paternal allele in *m-z+* embryos may compensate for the maternal loss (**Fig. S2B**). All RNA-seq biological replicates were highly reproducible (**Fig. S2A-B**), and the RNA-seq genome browser views confirmed the success of Cre-mediated depletion of *Dppa2* and *Dppa4* in the *m-z+* late 1-cell (**Fig. 4A**) and *m-z−* late 2-cell embryos (**Fig. 5A**).

**Figure 4.**
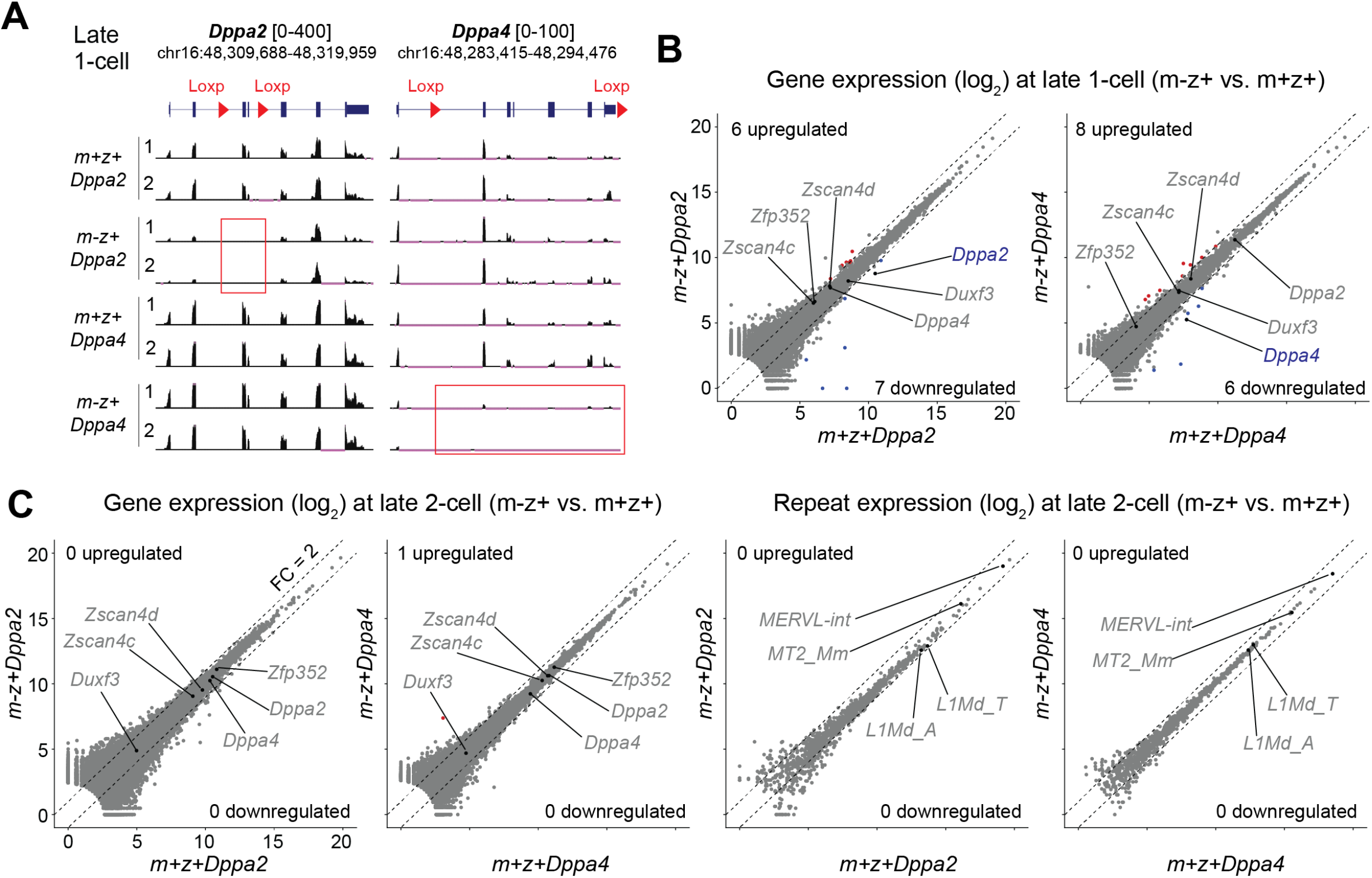
Maternal DPPA2 and DPPA4 are not responsible for activating *Dux* and ZGA. **A)** Genome browser views of indicated RNA-seq samples at *Dppa2* and *Dppa4* loci. Cre-mediated deletion of exons are highlighted by red boxes. **B)** Scatter plots comparing gene expression levels of 1-cell embryos (*m-z+* vs. *m+z+*). The x and y axes are normalized read counts by DESeq2 (log2)(Love et al. 2014). Differential gene expression criteria were fold change (FC) > 2, adjusted P-value < 0.05 and FPKM > 1. Note that the dots outside the dashed FC lines were not classified as differentially expressed because of their large P-values and/or low FPKM (**Table S1**). **C)** Scatter plots comparing gene/repeat expression levels of 2-cell embryos (*m-z+* vs. *m+z+*).

**Figure 5.**
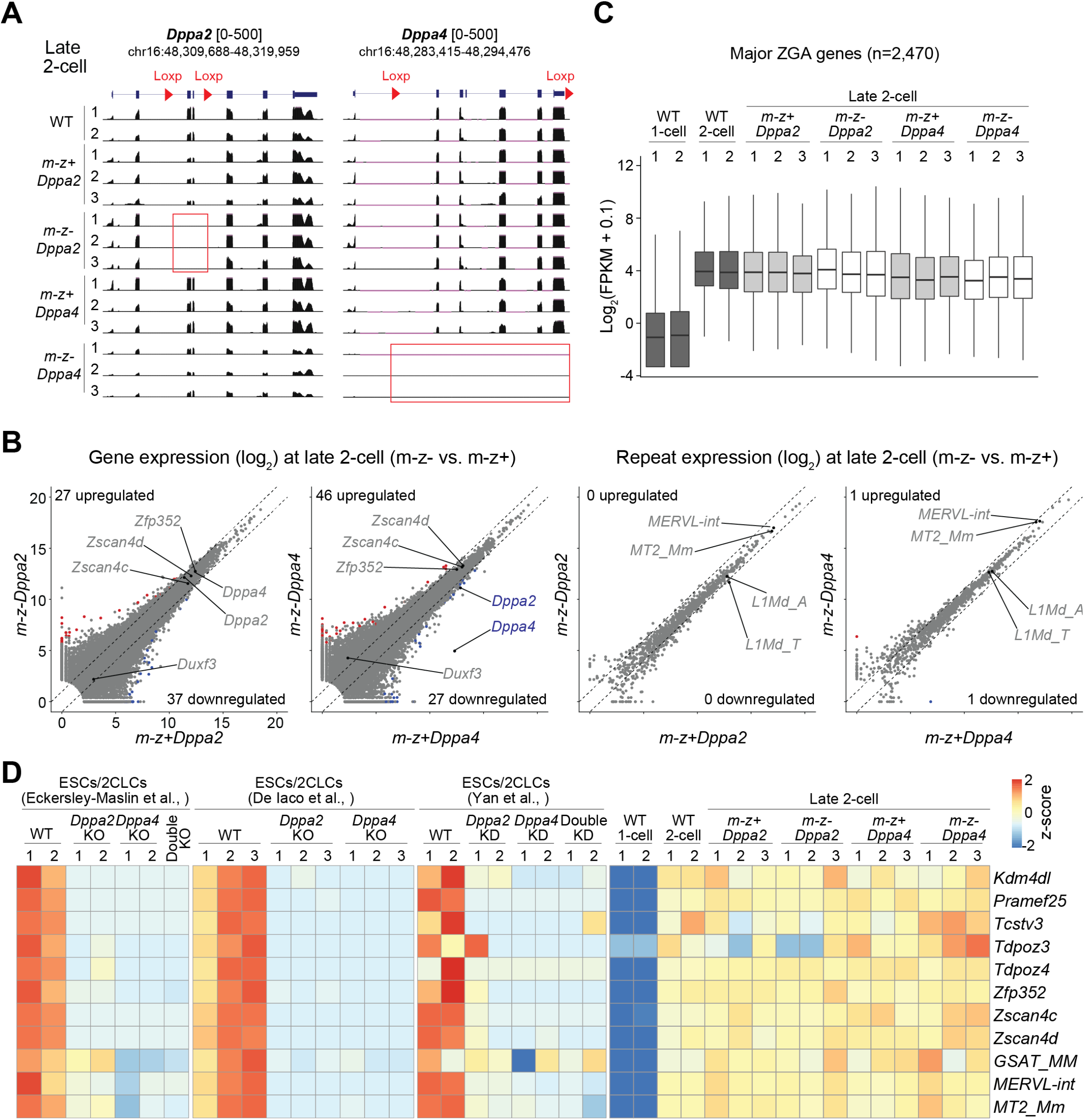
*Dppa2* and *Dppa4* maternal-zygotic KO embryos undergo normal ZGA. **A)** Genome browser views of indicated RNA-seq samples at *Dppa2* and *Dppa4* loci. Cre-mediated deletion of exons are highlighted by red boxes. **B)** Scatter plots comparing gene/repeat expression levels of 2-cell embryos (*m-z−* vs. *m-z+*). The x and y axes are normalized read counts by DESeq2 (log2)(Love et al. 2014). Differential gene expression criteria were fold change (FC) > 2, adjusted P-value < 0.05 and FPKM > 1. **C)** Boxplot illustrating the expression levels of major ZGA genes of indicated samples. The major ZGA genes were defined using cutoff: 2-cell/1-cell FC > 5, 2-cell FPKM > 3, adjusted p-value < 0.05. The middle lines represent medians. The box hinges indicate the 25^th^ and 75^th^ percentiles, and the whiskers indicate the hinger ± 1.5×interequatile range. **D)** Heatmap illustrating the expression levels of example genes/repeats in ESCs/2-cell like cells (2CLCs) and 2-cell embryos. The RNA-seq data of ESCs/2CLCs were from (De Iaco et al. 2019; Eckersley-Maslin et al. 2019; Yan et al. 2019).

We next performed comparative analyses in *m+z+*, *m-z+*, and *m-z−* embryos to identify differentially expressed genes (fragments per kilobase of transcript per million mapped reads (FPKM) >1, fold change (FC) >2, and adjusted *p*-value < 0.05). As expected, minimal changes in gene/repeat expression were observed in both *m-z+* and *m-z−* mutant embryos (**Fig. 4B-C, Fig. 5B, Table S1 and S2**). Note that both *Dux* and its target genes/repeats such as *Zscan4, Zfp352, MERVL-int*, and *MT2_Mm* were normally activated during minor and major ZGA. In addition, the young LINE-1 elements, including *L1Md_A* and *L1Md_T*, which are regulated by DPPA2 and DPPA4 independent of DUX in ESCs (De Iaco et al. 2019), were also normally expressed in late 2-cell embryos (**Fig. 4C, Fig. 5B**). Consistent with minimal transcriptome alterations, the major ZGA genes (n = 2470, 2-cell/1-cell: FC > 5, FPKM > 3, adjusted *p-*value < 0.05), including those previously reported to be regulated by DPPA2 and DPPA4 in ESCs, also showed normal activation (**Fig. 5C-D**). Thus, our data support that DPPA2 and DPPA4 are dispensable for *Dux* expression and ZGA in mouse early embryos.

Collectively, our data provide definite evidence that maternal DPPA2 and DPPA4 are not required to trigger the activation of *Dux* and other 2C embryo-specific genes during mouse ZGA. Although generation of double KOs were not feasible by natural mating due to their close genetic linkage, DPPA2 and DPPA4 should not compensate for each other for the following reasons. First, DPPA2 and DPPA4 function as a heterodimer (Nakamura et al. 2011; Hernandez et al. 2018). Loss of either protein causes comparable phenotypes to the double KOs during both ESC differentiation (*i.e.*, failure of developmental gene activation) (Eckersley-Maslin et al. 2020; Gretarsson and Hackett 2020) and embryogenesis (*i.e.*, lung developmental defects and perinatal lethality) (Madan et al. 2009; Nakamura et al. 2011). Second, our immunostaining analyses indicated that DPPA2 became almost undetectable when DPPA4 was depleted in early embryos (**Fig. 2C, Fig. 3C**), suggesting that similar results are expected from maternal-zygotic double KO. Therefore, compensation for each other should not explain the lack of apparent preimplantation phenotype for the DPPA2 and DPPA4 mutants analyzed in this study.

Together with the evidence that DUX does not initiate ZGA in embryos (Chen and Zhang 2019; Guo et al. 2019; De Iaco et al. 2020; Bosnakovski et al. 2021), this study further highlights the key differences between 2C-like cells and 2-cell embryos. In ESCs, both DUX and DPPA2/4 heterodimer are essential for establishing the 2C-like state (De Iaco et al. 2017; Hendrickson et al. 2017; De Iaco et al. 2019; Eckersley-Maslin et al. 2019; Yan et al. 2019). However, this is not the case in mouse embryos. Therefore, conclusions drawn from 2C-lilke cells should be carefully considered before being applied to the embryo scenario. Recently, TP53 has been identified as a maternal factor to regulate DUX and 2C-genes in both ESCs and embryos (Grow et al. 2021; Sun et al. 2021). Nonetheless, *Dux* also gets activated in *Tp53* maternal-zygotic KO embryos, although to a less extent than in WT (Grow et al. 2021). Thus, multiple pioneer factors may exist to trigger *Dux* and/or other minor ZGA genes (Kobayashi and Tachibana 2021).

Despite DPPA2 and DPPA4 are dispensable for ZGA, they may be required for maintaining a permissive chromatin state during gastrulation by counteracting DNA methylation, as suggested by the studies in ESCs (Eckersley-Maslin, 2020; Eckersley-Maslin et al., 2020; Gretarsson and Hackett, 2020). Loss of DPPA2 and DPPA4 may cause epigenomic defects around gastrulation, which ultimately contribute to the perinatal lethality phenotype (Madan et al., 2009; Nakamura et al., 2011). This hypothesis warrants to be examined by future studies.

## Materials and Methods

### Collection of mouse oocytes and preimplantation embryos

All animal studies were performed in accordance with guidelines of the Institutional Animal Care and Use Committee at Harvard Medical School. The procedures of GV and MII oocytes collection and *in vitro* fertilization (IVF) were described previously (Chen and Zhang 2019; Zhang et al. 2020).

### Generation of *Dppa2* and *Dppa4* mutant oocytes and embryos

The *Gdf9-Cre* transgenic line and the *Dppa4 fl* line were described previously (Lan et al. 2004; Nakamura et al. 2011). The *Dppa2 fl* allele was generated by the 2C-HR-CRISPR method (Gu et al. 2018). Specifically, *Cas9* mRNA (100 ng/μl), two sgRNAs (80 ng/μl each), and donor DNA PCR fragment (no biotin) (25 ng/μl) were co-injected into cytoplasm of each 2-cell blastomere using a Piezo impact-driven micromanipulator (Primer Tech). The PCR fragment with ~0.9-1kb homologous arms was used because high knock-in efficiency was reported for this donor DNA preparation method (Yao et al. 2018). After injection, the embryos were cultured for a few hours before transferred into oviducts of surrogate ICR strain mothers. The synthesis of *Cas9* mRNA and sgRNA was described previously (Wang et al. 2013). The donor DNA targeting vector was cloned using the Gibson Assembly method and the primers used for cloning are listed in **Table S3**. The *Dppa2 fl* F_0_ mice (BDF1 × BDF1) (Jackson 100006) were crossed with B6 (Jackson 000664) to confirm the germline transmission.

The breeding schemes were the same for *Dppa2* and *Dppa4* lines. Specifically, the *+/fl* lines were crossed with *Gdf9-Cre* to obtain *Gdf9-Cre, +/fl* males. The *Gdf9-Cre, +/fl* males were then crossed with *+/fl* females to obtain *Gdf9-Cre, fl/fl* males and *fl/fl* females. They were then crossed to generate *fl/fl* (control) and *Gdf-Cre, fl/fl* females (CKO) for experiments. The male mice that were heterozygous for Cre-mediated depletions were obtained by crossing *Gdf9-Cre, +/fl* females with WT B6 males. For all mouse lines, the tail tips were used for genotyping using primers listed in **Table S3**.

### Whole mount immunostaining

The immunostaining, image acquisition, and analyses were the same as previously described (Inoue et al. 2018). The primary and secondary antibodies are listed in **Table S3**.

### RNA-seq libraries preparation and data processing

The reverse-stranded total RNA-seq libraries (**Fig. 4** and **Fig. S2A**) were prepared using the SMARTer-Seq Stranded kit (Takara) following the manufacturer’s instructions. For single embryo RNA-seq (**Fig. 5** and **Fig. S2B**), the cDNA was synthesized using the SMARTer Ultra low Input RNA cDNA preparation kit (Takara). The cDNA was then used for genotyping by quantitative PCR and three embryos for each genotype (*i.e., m-z+* or *m-z−*) were selected for library construction using the Nextera XT DNA Library Preparation Kit (Illumina). The single embryo RNA-seq libraries were non-stranded and only PolyA+ RNA was captured. For all RNA-seq libraries, paired-end 75-bp sequencing was performed on a NextSeq 550 sequencer (Illumina). A summary of the generated data sets is available in **Table S2**.

The total RNA-seq (**Fig. 4**) and polyA RNA-seq (**Fig. 5**) were processed following the pipelines as previously described (Chen and Zhang 2019; Chen et al. 2021). The RNA-seq reads were mapped to mm10 assembly. RNA-seq pipeline and data processing R-codes are available at Github (https://github.com/YiZhang-lab/Nonessential_role_of_Dppa2_4_in_ZGA)

### Statistical analyses and data visualization

Statistical analyses were performed in R (www.r-project.org/). All sequencing tracks were visualized using the UCSC genome browser (Kent et al. 2002).

## Data availability

The RNA-seq data generated in this study have been deposited in the Gene Expression Omnibus under accession number GSEXXXXX (reviewer token: XXXXXXXX). The RNA-seq data of mouse oocytes and early embryos (**Fig. 1A**) were from GSE66582 (Wu et al. 2016) and GSE76505 (Zhang et al. 2018). The RNA-seq data of ESCs presented in **Fig. 5** were from GSE120952 (Eckersley-Maslin et al. 2019), GSE126621 (De Iaco et al. 2019), and GSE127811 (Yan et al. 2019).

## AUTHOR CONTRIBUTIONS

Y.Z. conceived the project; Z.C and Y.Z. designed the experiments; Z.C performed the experiments and analyzed the data sets; Z.X generated the *Dppa2* flox allele; Z.C and Y.Z. interpreted the data and wrote the manuscript.

## ACKNOWLEDGMENTS

We would like to thank Dr. Yota Hagihara for his help in some immunostaining experiments; Drs. Chunxia Zhang, Wenhao Zhang, Yota Hagihara, and Cheng-Jie Zhou for critical reading of the manuscript. This project was supported by NIH (R01HD092465) and HHMI. Y.Z. is an Investigator of the Howard Hughes Medical Institute.

## AUTHOR INFORMATION

The authors declare no competing financial interests.

## Supplemental figure legends

**Figure S1.**
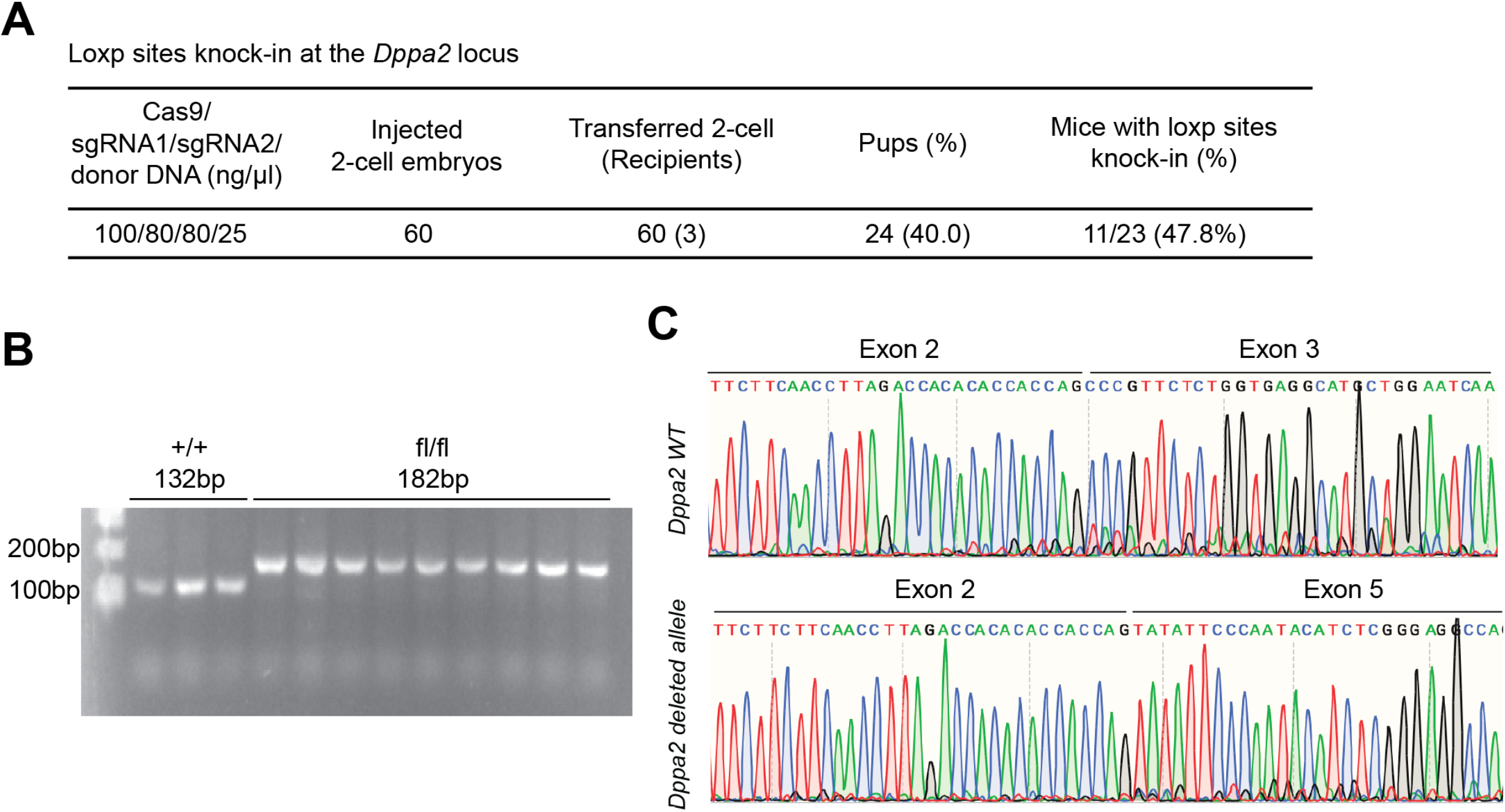
Generation of *Dppa2* flox allele and CKO model. **A)** Summary of the microinjection and embryo transfer experiments. **B)** Representative gel image for *Dppa2* flox allele genotyping. **C)** Sanger sequencing data showing the Cre-mediated deletion of exon 3-4 of *Dppa2* in oocytes.

**Figure S2.**
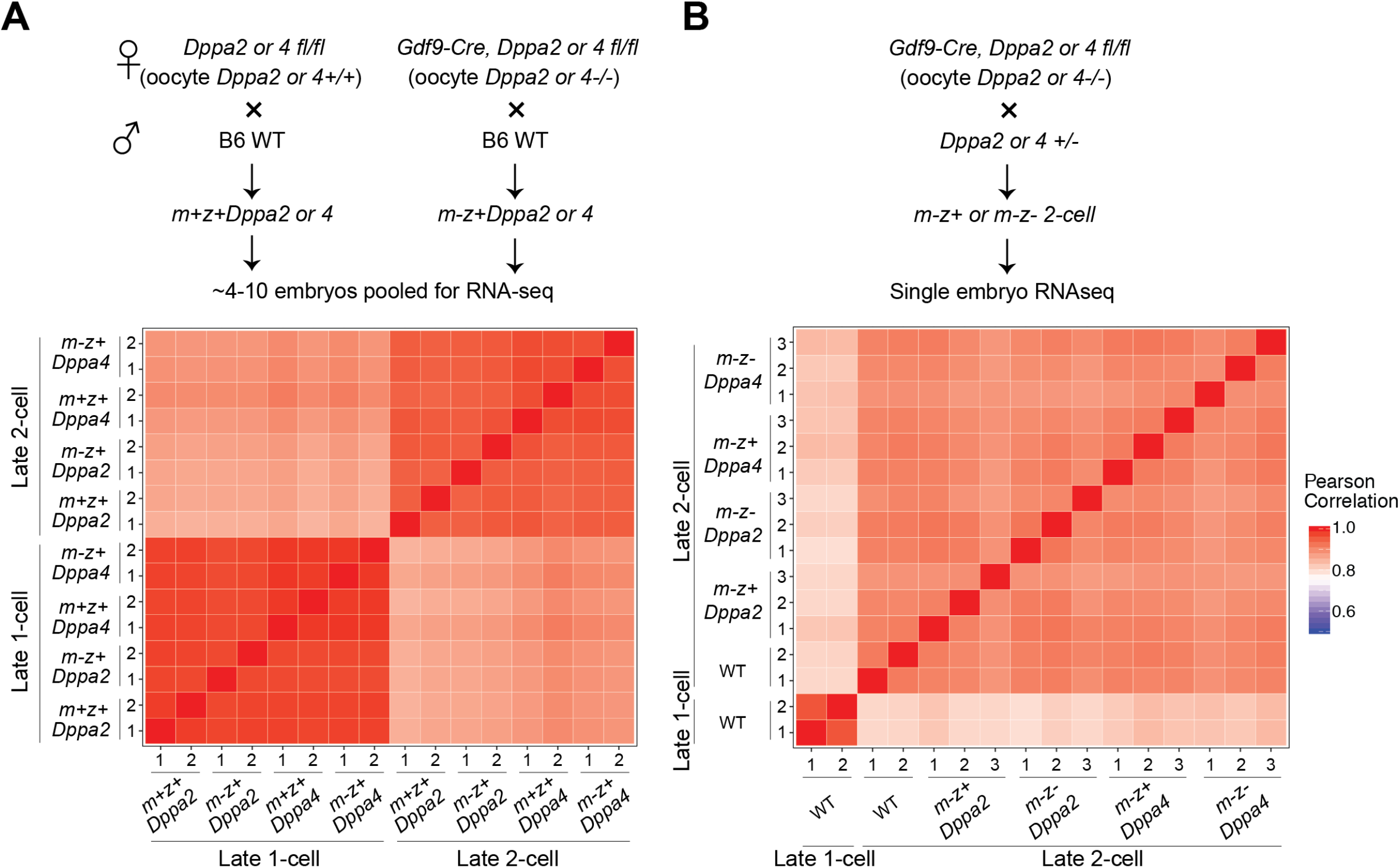
Reproducibility of RNA-seq experiments. For both panels **A-B)**, top panels showing the scheme for collecting pooled or single embryos for RNA-seq and bottom panels showing the Pearson correlation heatmaps.

## List of supplemental tables

**Supplemental table 1**: Differential gene expression analyses in *Dppa2* and *Dppa4* mutant embryos

**Supplemental table 2**: Summary of generated RNA-seq data sets

**Supplemental table 3**: List of primers and antibodies

## Notes

### Competing Interest Statement

The authors have declared no competing interest.

